# Epstein-Barr virus tegument protein BGLF2 in exosomes released from virus-producer cells assists *de novo* infection

**DOI:** 10.1101/2020.06.22.166280

**Authors:** Yoshitaka Sato, Masahiro Yaguchi, Yusuke Okuno, Hanako Ishimaru, Somi Ozaki, Takeshi Suzuki, Tomoki Inagaki, Miki Umeda, Takahiro Watanabe, Masahiro Fujimuro, Takayuki Murata, Hiroshi Kimura

**Author notes:** These authors contributed equally to this study (Yoshitaka Sato, Masahiro Yaguchi). Author order was determined alphabetically. To whom correspondence should be addressed: Yoshitaka Sato and Hiroshi Kimura.

## Abstract

Viruses must adapt to the environment of their host cells to establish infection and persist. Diverse mammalian cells, including virus-infected cells, secrete extracellular vesicles such as exosomes containing proteins and miRNAs, and use these vesicles to mediate intercellular communications. However, the roles of exosomes in viral infection remain unclear. Here we screened viral proteins to identify those responsible for the exosome-mediated upregulation of Epstein-Barr virus (EBV) infection. We found BGLF2 protein encapsulated in exosomes, which were released from EBV-infected cells. BGLF2 protein is a tegument protein that exists the space between the envelope and the nucleocapsid, and it is released into the cytoplasm shortly after infection. BGLF2 protein-containing exosomes enhanced viral gene expression and repressed innate immunity, thereby assisting the EBV infection. In summary, the EBV tegument protein BGLF2 is encapsulated not only encapsulated in viral particles, but also in exosomes secreted from infected cells. Therefore, BGLF2 may play a crucial role in establishing EBV latent infection.

## Introduction

Epstein-Barr virus (EBV), a member of the herpesviruses, mainly infects B cells. EBV infects about 90% of the world’s population asymptomatically, but can cause infectious mononucleosis and malignancies such as Burkitt lymphoma, T/NK cell lymphoma, nasopharyngeal carcinoma, and gastric carcinoma (Cohen et al., 2011; Young and Rickinson, 2004). EBV demonstrates a biphasic life cycle that includes a latent phase and a lytic (productive replication) phase (Murata and Tsurumi, 2014). After primary infection, although EBV preferentially remains latent in host cells without viral production, a limited number of viral genes are expressed. Under certain circumstances, latency is disrupted, and the virus shifts into the lytic phase. During this productive replication phase, all >80 viral genes are expressed, the viral genome undergoes replication, and progeny virions are produced (McKenzie and El-Guindy, 2015; Murata, 2014).

Similar to that of other herpesviruses, the EBV virion is composed of a viral genomic DNA-containing nucleocapsid surrounded by a viral envelope with glycoprotein spikes on its surface (Diefenbach, 2015; Knipe et al., 2007). Located between the nucleocapsid and the outer viral envelope is the viral tegument which includes >10 viral proteins called tegument proteins (Johannsen et al., 2004) that are involved not only in morphogenesis, envelopment, and egress of progeny virions but also in primary *de novo* infection (Sathish et al., 2012). Because they are released shortly after infection into the cytoplasm, tegument proteins play major roles in infectivity by modulating the intracellular environment. Indeed, we recently showed that the EBV tegument protein BGLF2 increases viral infectivity (Konishi et al., 2018).

Extracellular vesicles such as exosomes are secreted from eukaryotic cells. These vesicles can contain proteins, RNA, microRNA, or DNA, and they serve as crucial mediators of intercellular communication involved in organismal homeostasis and disease states (Raab-Traub and Dittmer, 2017; Record et al., 2014; Yanez-Mo et al., 2015). The roles of extracellular vesicles have received considerable attention in recent years. Exosomes secreted by virus-infected cells contain viral materials (Meckes and Raab-Traub, 2011). Interestingly, EBV-infected cell lines secrete a higher number of extracellular vesicles than uninfected cells (Hurwitz et al., 2017), leading us to hypothesize that EBV utilizes exosomes to influence infection and pathogenesis by facilitating viral replication or activating immune evasion strategies. For instance, the viral oncoprotein latent membrane protein 1 (LMP1) has been detected in exosomes (Houali et al., 2007; Sato et al., 2017; Vazirabadi et al., 2003). LMP1-positive exosomes activate the extracellular signal-regulated kinase and phosphoinositide 3- kinase (PI3K)/Akt signaling pathways in the recipient cells (Meckes et al., 2010).

Although EBV encodes more than 80 genes, many of which are lytic proteins, the majority of viral proteins detected in exosomes are latent proteins (Zhao et al., 2019), because these latent proteins have been identified using a forward genetics approach. Therefore, it is still unclear which EBV lytic proteins are encapsulated in exosomes and function during infection.

In this study, we use a reverse genetics approach to investigate the role of EBV protein-containing exosomes during infection. We performed a series of screenings and identified the tegument protein BGLF2, which enhanced *de novo* infection via exosomes. Our results suggest that this tegument protein functions via extracellular vesicle transfer like that of virus particles, ensuring efficient and robust modulation of the microenvironment for the infection.

## Results

### Screening for EBV proteins that modulate infection via exosomal transfer

To explore the roles of exosomes containing EBV proteins during infection, we generated an exosome library, from 293T cells expressing 71 different epitope-tagged EBV lytic proteins (Konishi et al., 2018), and screened for EBV genes that both modulated infection and were detected in exosomes (Figure 1A). EBV-negative Akata(-) B-cells treated with each exosome from the library were experimentally infected with EBV, and the infectivity at 48 h post-infection (hpi) was compared (Figure 1B, upper panel). Additionally, the exosome library was subjected to immunoblot analysis with an anti-hemagglutinin (HA) antibody to detect EBV proteins in the exosomes (Figure 1B, lower panel). By integrating multiple screenings, we found BGLF2 and BLRF1 within exosomes, and that they enhanced EBV infection. Because BLRF1 encodes envelope glycoprotein N, which localized to the membrane, we here focus on the tegument protein BGLF2.

**Figure 1:**
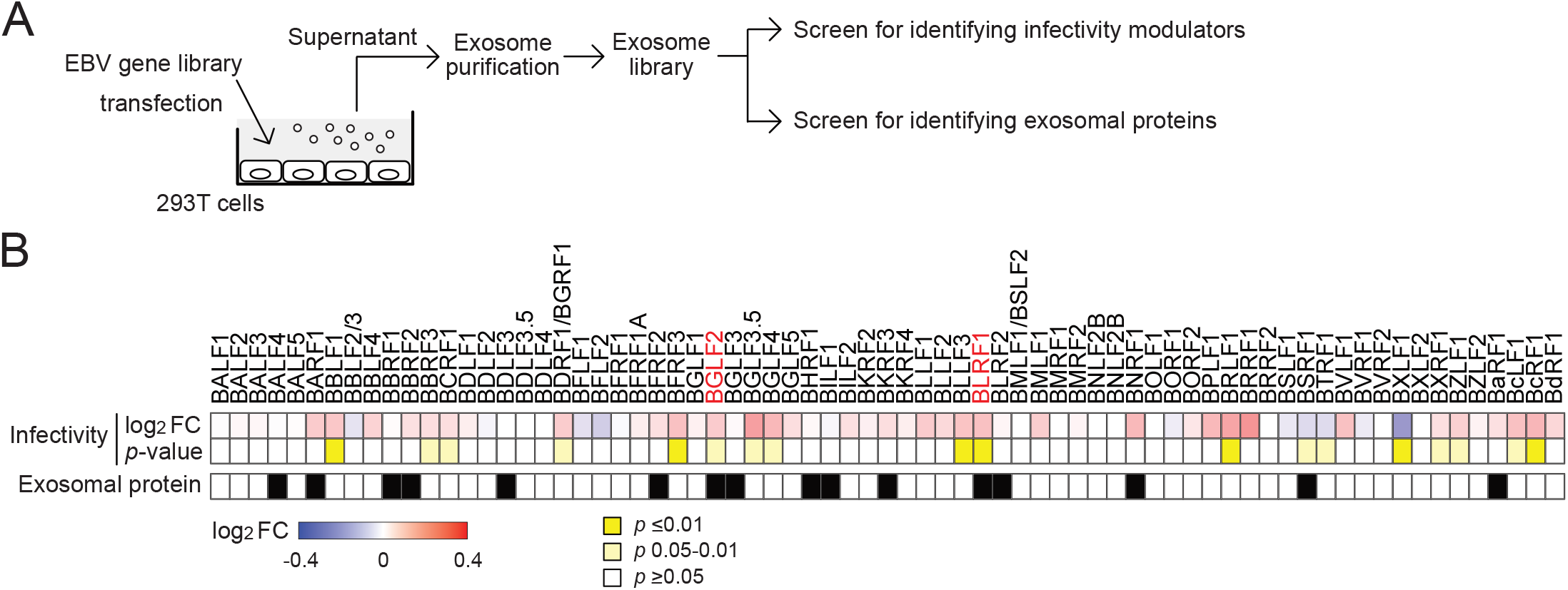
Combined screenings for the viral proteins that regulate the infectivity via exosomal transfer. (A) The workflow of the screenings. Exosomes were purified from the conditioned media of HEK293T cells that were transfected with each plasmid expressing an EBV gene. For analyzing modulators of infectivity, EBV-negative Akata(-) cells were pre- treated with purified exosomes and then infected with EGFP-EBV. The infectivity was determined at 48 hpi by FACS analysis. For identifying exosomal proteins, purified exosomes were analyzed by immunoblot with an anti-HA antibody. (B) Heatmap showing fold change (FC) of infectivity of EBV with Akata(-) cells in the presence of each exosome at 48 hpi. Infectivity changes (top panel) reflect the log^2^ FC relative to the control exosome purified from empty vector-transfected cells. p values are shown on the middle panel. The viral protein in purified exosomes is indicated as black on the bottom panel.

### Exosome-mediated release of BGLF2 in culture supernatant

Because exosomes were purified by a precipitation-based method in our screening for easy handling, we could not rule out the possibility of non-exosome contamination (Patel et al., 2019). To examine the incorporation of BGLF2 in the exosomes, we isolated exosomes, using an affinity-based method, from the culture supernatant of 293T cells expressing BGLF2-HA. BGLF2 was also detected in the exosomes purified by both exosome-affinity resin (Figure 2A) and anti-CD9 antibody (data not shown). Size range and morphology of exosomes were characterized via nanoparticle tracking analysis (NTA) and transmission electron microscopy (TEM), respectively (Figure 2B and C). We confirmed that cellular cytochrome C (Yoshioka et al., 2013) could not be detected in the purified exosomes, suggesting no contamination of any other types (Figure 2D). Moreover, we confirmed that the detergent treatment of the culture supernatant impeded the BGLF2 detection in the exosomes, suggesting that BGLF2 was encapsulated within the exosomes (Figure 2E).

**Figure 2:**
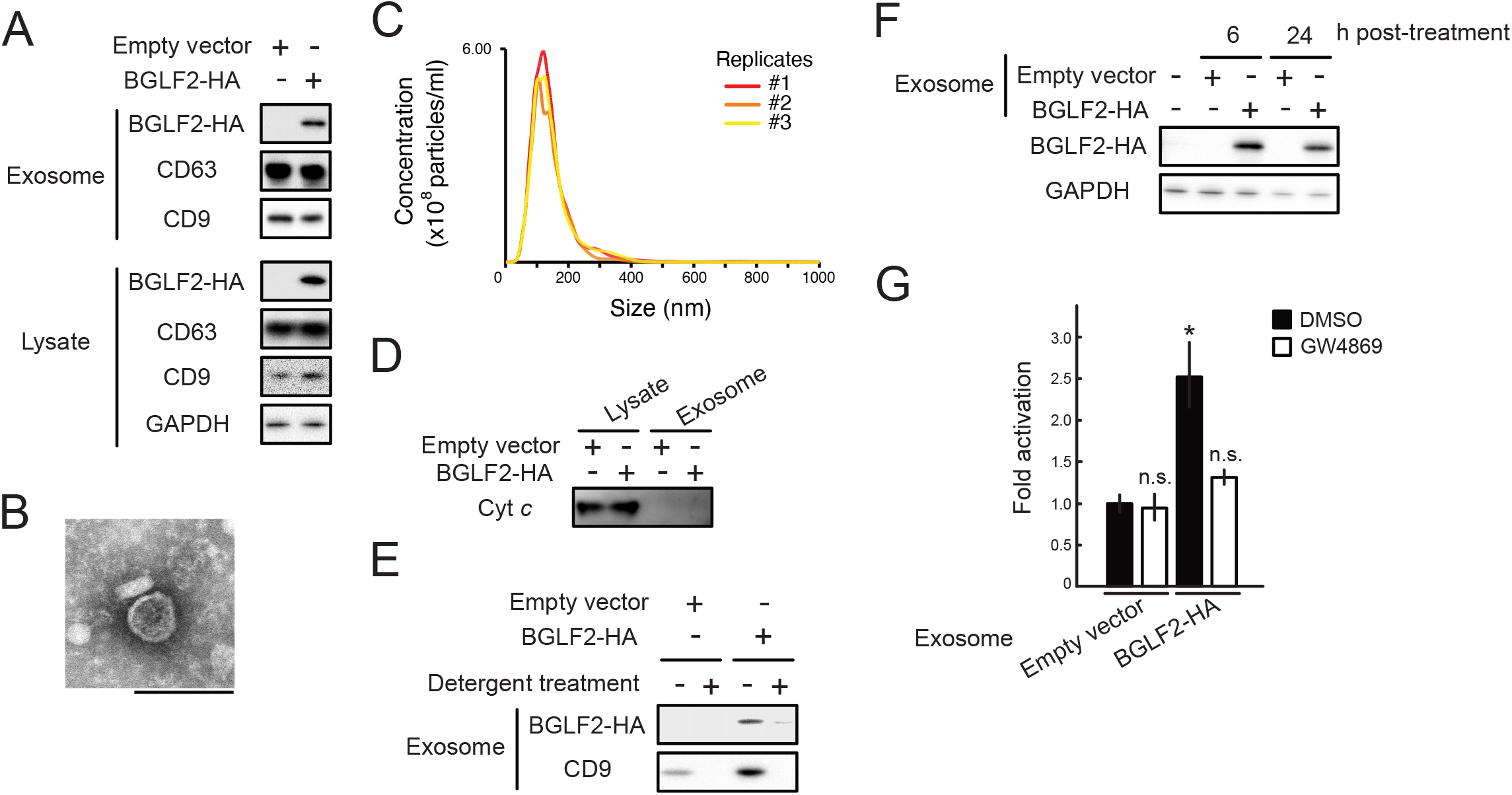
Functional BGLF2 proteins are transferred to the recipient cells via exosomes. (A) BGLF2 protein-containing exosomes were secreted from BGLF2-expressing cells. HEK293T cells were transfected with HA-tagged BGLF2-expression or empty plasmid. Exosomes were isolated from the supernatant of the cells. CD63 and CD9 served as exosome markers. (B) TEM image of the exosomes purified from BGLF2-expressing HEK293T cells. Scale bar represents 100 nm. (C) NanoSight profiles showing size distribution for exosomes purified from BGLF2- expressing HEK293T cells. (D) Western blot results showing the absence or underrepresentation of negative exosome marker cytochrome-C (Cyt *c*) in isolated exosomes. (E) BGLF2 protein was incorporated into exosomes. Exosomes were purified from supernatants treated with or without 0.25% Triton-X 100 for 5 min at room temperature. (F) BGLF2 was delivered to the recipient cells via exosomes. HEK293T cells were treated with BGLF2-containing exosomes for the indicated number of hours and then washed extensively and harvested. Cell lysates were applied to immunoblot analysis. (G) BGLF2 transferred via exosomes activated the AP-1 promoter. Exosomes were purified from HEK293T cells transfected with BGLF2-expression plasmid or empty plasmid in the presence or absence of GW4869. HEK293T cells transfected with the pAP-1-Luc and promoterless-Rluc plasmids, and then treatment with BGLF2- containing exosomes. Cell lysates were obtained at 24 h post-treatment and analyzed using luciferase reporter assays. Values (mean ± SEs) were calculated from three independent experiments. Asterisks, p<0.05; n.s., not significant.

BGLF2 stimulates the activator protein (AP)-1 signaling pathway (Konishi et al., 2018; Liu and Cohen, 2016). To assess the function of BGLF2-containing exosomes, we added them to cells that were transfected with a pAP-1-Luc reporter and control plasmids. As shown in Figure 2F, BGLF2 was taken up by the recipient cells, and treatment with BGLF2-containing exosomes enhanced AP-1 reporter activity (Figure 2G). GW4869, a compound blocking exosome release by neutral sphingomyelinase inhibition (Trajkovic et al., 2008), prevented AP-1 activation (Figure 2G). These findings indicate that functional BGLF2 is transferred via exosomes to recipient cells.

We further evaluated whether BGLF2-containing exosomes could be released from infected cells. Conditioned medium from lytic-induced Akata/EBV-EGFP cells by IgG was subjected to floating density gradient ultracentrifugation and separated into 18 fractions (Figure 3A). Each fraction was characterized by immunoblotting with exosome marker antibodies and quantitative polymerase chain reaction (qPCR) analysis for EBV genome detection (Figure 3B). Three fractions containing exosomes (E), virions (V), and control (C) were pooled and further evaluated as the antibody against BGLF2 did not exhibit high sensitivity. As shown in Figures 3C and D, BGLF2 proteins were detected in pool E, which was free of viral genome and viral capsid antigen (VCA), indicating that infected cells released BGLF2-containing exosomes during the lytic replication.

**Figure 3:**
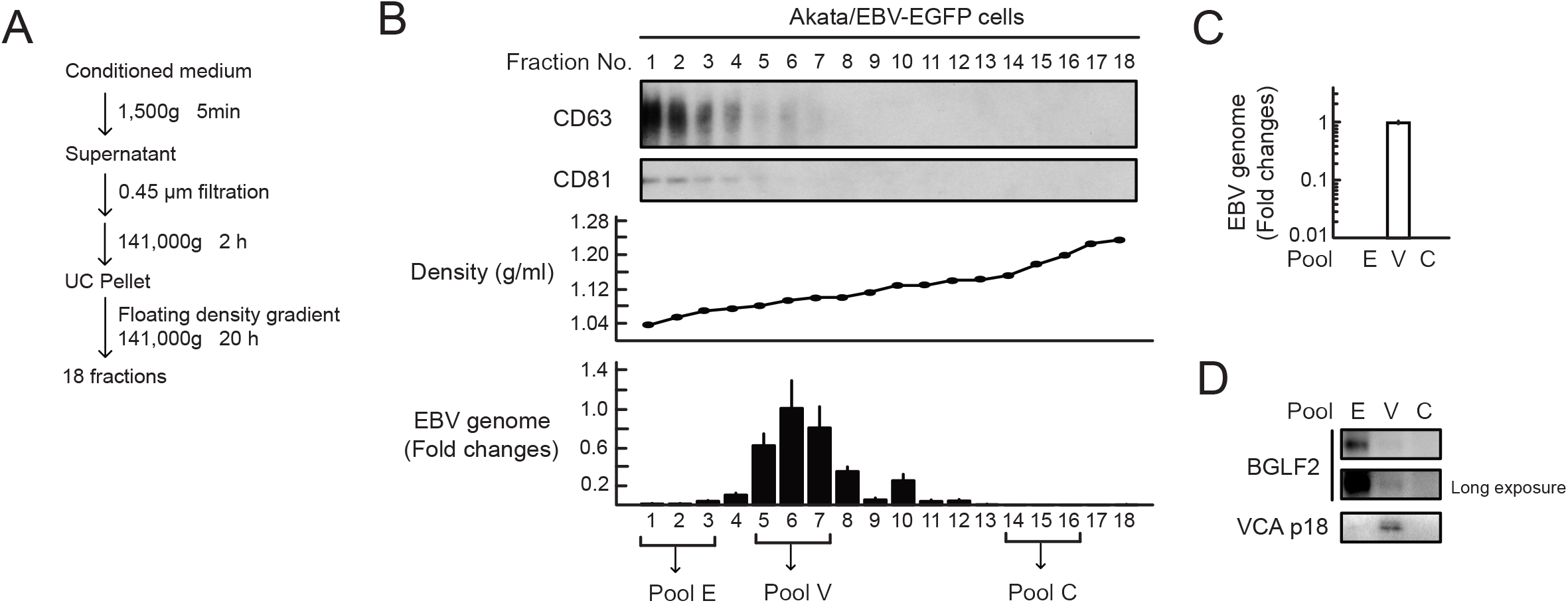
BGLF2-containing exosomes are released from infected cells. (A) Centrifugation protocol and workflow for separation and enrichment of exosomes and virions. (B) Exosome marker proteins were enriched in fractions #1-3 after iodixanol floating density gradient centrifugation. For the Western blotting analysis, proteins in each fraction were concentrated by acetone precipitation. CD63 and CD81 served as exosome markers. The EBV genome in each fraction was quantified by qPCR. (C) and (D) Pool E (fraction #1-3) containing the BGLF2 protein. Each pool (E, fraction #1-3; V, fractions #5-6; and C, fractions #14-16) was analyzed by qPCR (C) and Western blot (D). Data are presented as the means ± SE. Viral capsid antigen (VCA) p18 served as virion marker.

### EBV BGLF2 orthologs of in herpes simplex virus-1 and human cytomegalovirus are incorporated into exosomes

To define the regions of BGLF2 responsible for packaging into exosomes, we created truncation mutants of BGLF2, which is 336 amino acids (aa) in length, and then tested each for its incorporation (Supplementary Figure S1A). Bortezomib, a proteasome inhibitor, was used for stabilizing deletion mutants. As shown in Figure S1A, the deletion of 98-158 aa precluded the incorporation of BGLF2 into exosomes, although deletion of the C-terminal domain did not. Thus, the incorporation of BGLF2 into exosomes required its central domain. In line with previous reports (Cai et al., 2017; Salsman et al., 2008), BGLF2 exhibited a pan-cellular localization pattern without a restricted colocalization with CD63, a marker for multi-vesicular bodies (Supplementary Figure S1B).

All herpesviruses possess a BGLF2 homologue (Liu and Cohen, 2016). Therefore, we tested whether the orthologs of EBV BGLF2 such as herpes simplex virus type 1 (HSV-1) UL16 and human cytomegalovirus (HCMV) UL94 proteins were incorporated into exosomes. HSV-1 UL16 and HCMV UL94 proteins were detected in exosomes purified from cells expressing HSV-1 UL16 and HCMV UL94 (Figure 4). These findings indicate a conserved role of exosomes containing BGLF2 or its herpesviral orthologs proteins.

**Figure 4:**
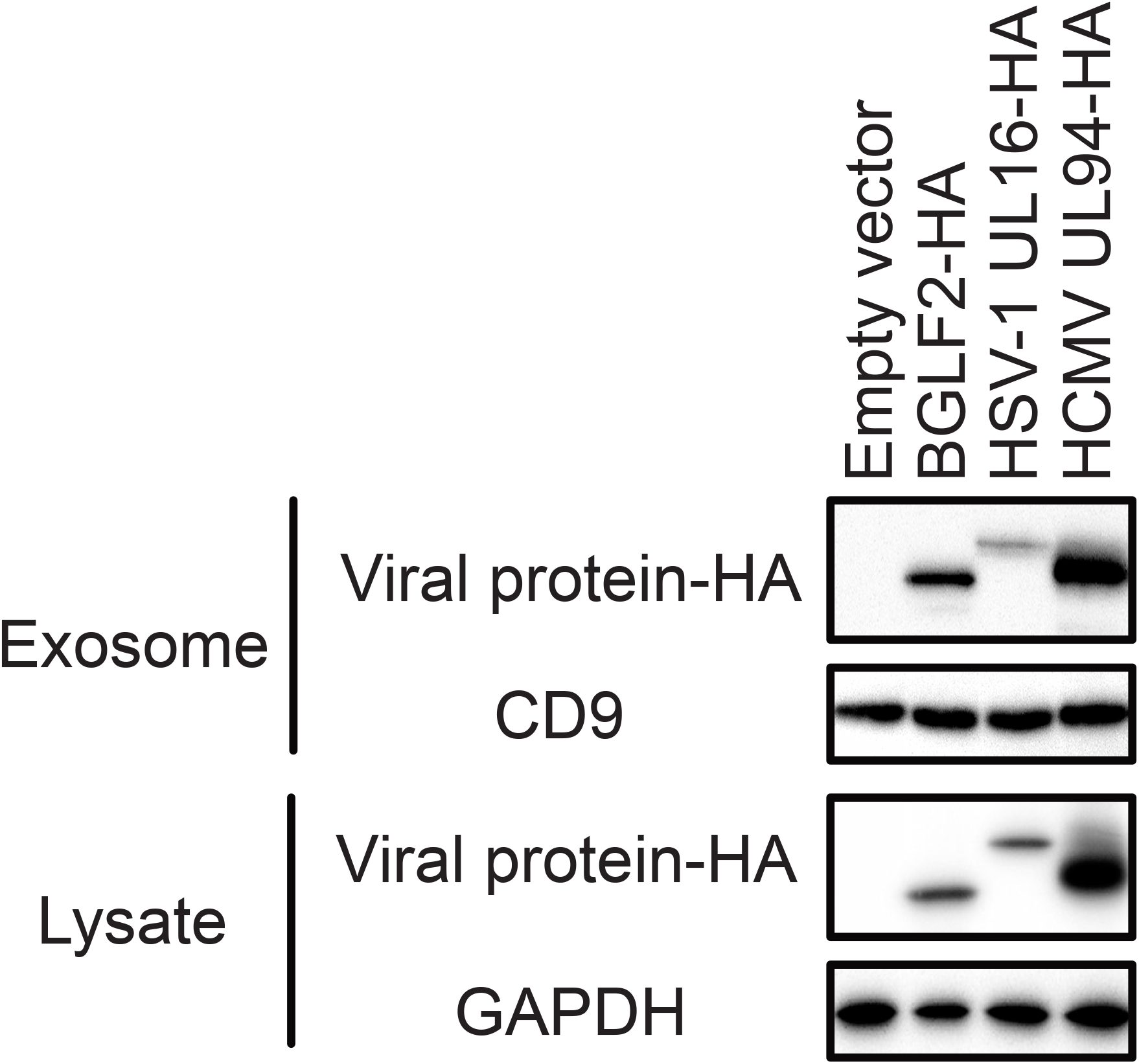
The orthologs of BGLF2 are incorporated into exosomes. HEK293T cells were transfected with expression plasmids containing HA-tagged BGLF2 or its HSV-1 or HCMV orthologs. Exosomes were isolated from the supernatant of the cells.

### Extra-virion particle BGLF2 enhances the infectivity of BGLF2-KO EBV

Next, we examined whether exosomal-transferred BGLF2 supports infection of using a BGLF2-knockout (BGLF2-KO) EBV. The knockout of BGLF2 reduces EBV infectivity upon *de novo* infection of B-cells (Konishi et al., 2018). Treatment with exosomes containing BGLF2 significantly enhanced the infectivity of BGLF2-KO EBV compared with a control treatment (Figure 5A).

**Figure 5:**
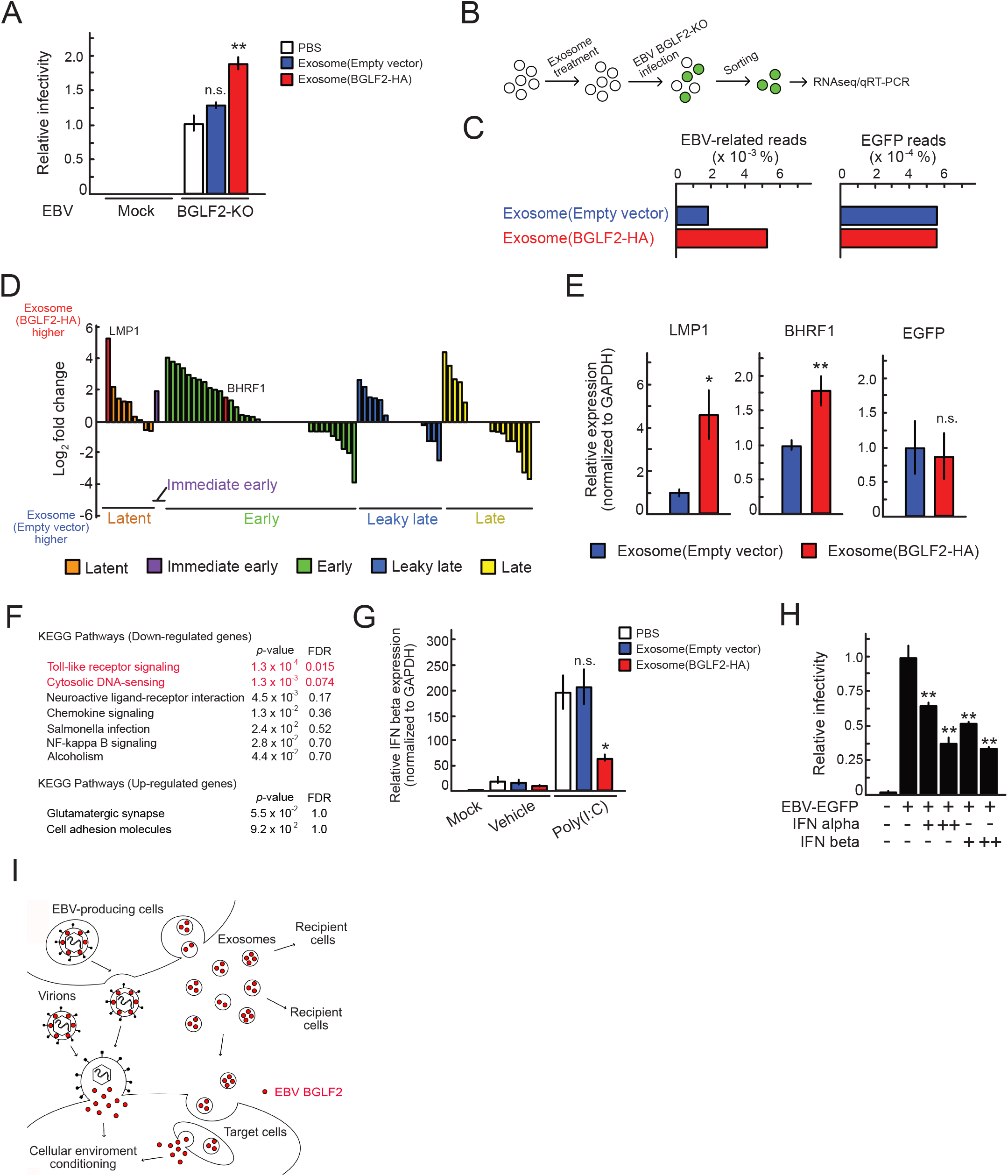
BGLF2-containing exosomes enhance the infectivity of BGLF2-KO EBV. (A) Akata(-) cells were infected with BGLF2-KO EBV in the presence of BGLF2- containing exosomes. After two days, GFP positivity was determined by FACS. Results are presented as means ± SE of at least three independent experiments and are shown as relative infectivity to control treatment with PBS (infectivity value of 1). Double asterisks, p<0.01; n.s., not significant. (B) The workflow of sample collection for RNA-seq/qRT-PCR analyses. Akata(-) cells were pre-treated with exosomes for 16 h, and then infected with BGLF2-KO EBV. The infected cells were collected via FACS at 24 hpi. Total RNA extracted from the sorted cells was subjected to RNA-seq and qRT-PCR. (C) Gene expression levels were normalized as counts per million followed by log2- transformation with a pseudo-count of 1. Each bar indicates the log2 fold change of a viral gene expression between treatment with exosomes with or without BGLF2. Viral gene expression kinetics are categorized into five groups: latent, immediate early, early, leaky late, and late (Djavadian et al., 2018). (D) Read counts of EBV genes in BGLF2-KO EBV-infected cells that were treated with BGLF2-containing or control exosomes by RNA-seq. (E) Validation of enhanced EBV gene expression by qPCR. Data are presented as means ± SE. Samples were tested in duplicate. Asterisk, p<0.05; double asterisks, p<0.01; n.s., not significant. (F) DAVID analysis of RNA-seq data from BGLF2-KO EBV-positive cells treated with BGLF2-containing or control exosomes. FDR; false discovery rate. (G) HEK293 cells were treated with BGLF2-containing exosomes for 16 h. Cells were challenged with transfection of poly(I:C) (2 μg/ml) for 2.5 h. RT-qPCR was conducted. Data are presented as the means ± SE from three independent experiments. Asterisk, p<0.05; n.s., not significant. (H) Type I IFN inhibited EBV infection. Akata(-) cells were infected with EBV-EGFP in the presence of IFN alpha and beta (100 or 500 Unit/mL). After two days, GFP positivity was determined by FACS. Results are presented as means ± SE of three independent experiments. Double asterisks, p<0.01; n.s., not significant. (I) BGLF2 protein in exosomes cooperates with the tegument protein in the virions to optimize the cellular condition for EBV infection. The tegument protein BGLF2 is incorporated into both exosomes and virions. BGLF2-containing exosomes are released from infected cells and transferred to the recipient cells.

To evaluate the effect of BGLF2-containing exosomes on EBV infection, we performed RNA-seq analysis (Figure 5B). Akata(-) cells were pre-treated with BGLF2- containing exosomes or control exosomes and then infected with BGLF2-KO EBV. We collected infected cells by fluorescence-activated cell sorting (FACS) at 24 hpi according to EGFP-positivity, and performed RNA-seq to obtain transcriptome information. As shown in Figure 5C and 5D, treatment with BGLF2-containing exosomes induced expression of EBV genes, although the expression level of EGFP, which was driven from a eukaryotic promoter (Delecluse et al., 1998), was the same. This enhanced EBV mRNA expression was validated using qPCR (Figure 5E). LMP1 was one of the most significantly expressed viral genes between the groups treated with exosomes with or without BGLF2 (Figure 5C).

Among down-regulated genes following treatment with BGLF2-containing exosomes, the terms “Toll-like receptor signaling” and “cytosolic DNA-sensing” were enriched by DAVID analysis (Huang da et al., 2009) of RNA-seq data (Figure 5F). This is consistent with a recent report that ORF33, the ortholog of BGLF2 in Kaposi’s sarcoma-associated herpesvirus (KSHV), suppresses the interferon (IFN) beta production pathway for immune evasion (Yu et al., 2020). Consequently, we investigated whether BGLF2-containing exosomes modulated the innate immune response. Polyinosinic-polycytidylic acid [poly(I:C)] was used to stimulate the TLR3- mediated IFN beta production pathway (Takeuchi and Akira, 2010). As shown in Figure 5G, BGLF2-containing exosomes inhibited IFN beta mRNA expression under poly(I:C) stimulation. We confirmed that type I IFNs inhibited EBV infection in a dose-dependent manner (Figure 5H).

Together, our data indicate that BGLF2 is encapsulated not only encapsulated in viral particles, but also in exosomes secreted from infected cells. BGLF2 delivered via exosomes promotes optimal EBV infection by enhancing viral gene expression and repressing innate immunity (Figure 5I).

## Discussion

Virally infected cells release virions as infectious particles and non-infectious particles. All herpesvirus-infected cells are known to release L-particles in addition to virions (Chang et al., 1997; Granato et al., 2008; Irmiere and Gibson, 1983). L-particles are non-infectious because they are composed of the virus envelope and tegument proteins but lack viral genome and nucleocapsid proteins. This conserved release of L-particles may shed light on an essential role of non-infectious particles in creating a microenvironment that facilitates their replication, spread, and persistence in the host cells. Recently, it was shown that L-particles mediate immune evasion by CD83 down-modulation on mature dendritic cells (Heilingloh et al., 2015).

BGLF2 protein impairs type I IFN signaling through Tyk2 (Liu et al., 2020) and also dampens NF-κB activity (Chen et al., 2019) in a cell-dependent manner during lytic infection. Expression of KSHV ORF33 (the ortholog of BGLF2) suppresses the IFN beta production pathway (Yu et al., 2020). Here, we demonstrated that exsosomal tegument protein BGLF2 was transferred to the target cells and in turn enhanced EBV infectivity by manipulating the cellular environment (Figure 5I). BGLF2 delivered via exosomes increase the expression of EBV genes (Figure 5C-E). The LMP1 expression is upregulated by the binding of AP-1 with CCAAT enhancer binding proteins (C/EBPs) on its promoter (Noda et al., 2011). The EBV gene BHRF1 is highly expressed during the first two days after infection (Mrozek-Gorska et al., 2019; Wang et al., 2019). These findings imply a role for BGLF2 in viral gene expression in infected cells, consistent with previous reports that BGLF2 activates p38, JNK, and AP-1 to induce viral gene expression (Konishi et al., 2018; Liu and Cohen, 2016). Furthermore, exosomes prepared from BGLF2-expressing 293T cells inhibited IFN beta production stimulation (Figure 5G), protecting infected cells from a type I IFN-mediated response during *de novo* infection. Exosomal BGLF2 protein was active in the recipient cells (Figure 2G and 4) suggesting that exosomes serve as a carrier of herpesvirus tegument proteins to target cells. Indeed, BGLF2-containing exosomes were released from EBV-infected cells that lytically produced infectious virions (Figure 3).

EBV can hijack and use extracellular vesicles such as exosomes. It is known that EBV increases the extracellular vesicle secretion (Hurwitz et al., 2017) and EBV miRNAs delivered via exosomes increase the severity of EBV-mediated lymphoproliferative disease (Higuchi et al., 2018). Although exosomes are released in exocytic bursts upon the fusion of the multivesicular bodies to the cell membrane (Colombo et al., 2014), the final envelopment site for EBV is the Golgi apparatus (Nanbo et al., 2018), indicating that these non-infectious particles (L-particles and exosomes) are generated in non-Golgi compartments. Furthermore, BGLF2 and its orthologs in HSV-1 and HCMV were incorporated into exosomes (Figure 4), indicating that manipulation of the cellular environment by exosomes is a conserved mechanism among herpesvirus. The results of this study also suggest a previously unforeseen essential role for non-infectious particles in viral protection.

## Materials and Methods

### Cells, plasmids and reagents

HEK293T (ATCC CRL-3216) and HEK293/EBV(dBGLF2) cells (Konishi et al., 2018) were maintained in DMEM supplemented with 10% fetal bovine serum (FBS). Akata(–) (Shimizu et al., 1994), and Akata/EBV-EGFP (Masud et al., 2017) cells were cultured in RPMI1640 medium containing 10% FBS. AGS/EBV-EGFP cells (a kind gift from Hironori Yoshiyama) (Katsumura et al., 2009) were grown in F-12 HAM’s medium supplemented with 10% FBS and 400 μg/mL G418.

The BZLF1-expression plasmid and HA-tagged EBV lytic protein-expression library were described previously (Konishi et al., 2018; Sato et al., 2009). For expression of C-terminal HA-tagged HSV-1 UL16 and HCMV UL94, constructs expressing UL16 and UL94 in pcDNA3 (Thermo Fisher Scientific, Waltham, MA, USA) were prepared by PCR and the In-Fusion cloning system (Takara Bio, Kusatsu, Shiga, Japan). The inserted DNA sequence of each vector was confirmed by direct DNA sequencing. The reporter plasmid pAP1-Luc and promoterless-RLuc (pGL4.70) were purchased from Promega (Madison, WI, USA).

Bortezomib (Cat# A10160; AdooQ BioScience, Irvine, CA, USA) was used at a concentration of 100 nM. Recombinant human type I IFN alpha (Cat# 11200-1) and IFN beta (Cat# 11415-1) were purchased from PBL Assay Science (Piscataway, NJ, USA). For inhibition of exosome secretion, cells were treated with 10 μM GW4869 (Cat# 13127; Cayman Chemical, Ann Arbor, MI, USA). Transient transfection of poly(I:C) (Cat# 4287/10; R&D Systems, Minneapolis, MN, USA) was performed with Lipofectamine 2000 (Thermo Fisher Scientific) according to the manufacturer’s protocol.

### Antibodies and immunoblotting

Anti-HA (3F10) rat antibody was purchased from Sigma-Aldrich (St. Louis, MO, USA). Anti-CD63 (Ts63) mouse and anti-VCA (PA1-73003) goat antibodies were obtained from Thermo Fisher Scientific. Rabbit anti-GAPDH (14C10), rabbit anti-CD9 (D8O1A), and horseradish peroxidase-conjugated secondary antibodies were purchased from Cell Signaling Technology (Danvers, MA, USA). Anti-CD81 (5A6) mouse antibody was obtained from Santa Cruz Biotechnology (Dallas, TX, USA). Antiserum against EBV BGLF2 was prepared by immunizing a rabbit with the synthetic peptide NH2-CAHVNILRGWTEDDSPGTS-COOH, and the serum was purified by affinity purification using the same peptide (Cosmo Bio, Tokyo, Japan) (Supplementary Figure S2). Immunoblot and signal detection were performed as described previously (Sato et al., 2009).

### Virus preparation

EGFP-EBV was obtained from the 8-day-old cell-free supernatant of AGS/EGFP-EBV cells. The cell-free supernatant was filtered through 0.45 μm filters and then used as a virus stock. For the preparation of the BGLF2-KO virus, HEK293/EBV(dBGLF2) cells (Konishi et al., 2018) were transfected with BZLF1 expression plasmid using Lipofectamine 2000 (Thermo Fisher Scientific) according to the manufacturer’s instructions. Cells and media were harvested and freeze-thawed, and cell debris was removed. The supernatant after centrifugation was filtered through 0.45 μm filters and then used as a virus stock. EBV-negative Akata(-) cells were infected with the virus, and EGFP-positive cells were counted by FACS to measure the viral titer.

### Exosome purification

Exosome-containing conditioned media were centrifuged at 2000 × g to remove cells. Exosomes were purified with a Total Exosome Isolation Kit (Thermo Fisher Scientific) for screenings or exoEasy Maxi Kit (Qiagen, Hilden, Germany) according to the manufacturer’s protocols.

Size and particle number were analyzed using the LM10 or NS300 nanoparticle characterization system (NanoSight; Malvern Panalytical, Worcestershire, United Kingdom).

### Transmission electron microscopy (TEM)

For negative staining TEM analysis, the samples were absorbed on formvar film coated copper grids and were stained with 2% phosphotungstic acid solution (pH 7.0) for 1 min. The grids were observed using a transmission microscope (JEM-1400Plus; JEOL Ltd., Tokyo, Japan) at an acceleration voltage of 100 kV. Digital images were captured using a CCD camera (EM-14830RUBY2; JEOL Ltd.).

### Floating density gradient ultracentrifugation

Semi-confluent Akata/EBV-eGFP cells were washed with PBS twice and suspended in medium containing EV-free FBS and anti-human IgG for the induction of lytic replication. After 72 h, 100 ml of the conditioned medium was centrifuged at 1,500 g and 4°C for 5 min to remove cells and debris. The supernatant was filtered by using a 0.45-μm filter (Merck, Darmstadt, Germany) and extracellular vesicles including exosomes and virions were pelleted by ultracentrifugation at 141,000 g at 4 °C for 2 h in a swinging-bucket rotor (SW28; Beckman Coulter, Brea, CA, USA). The pellet was resuspended in filtered PBS and centrifuged at 18,300 g and 4°C for 10 min to remove insoluble aggregates. The supernatant was mixed with iodixanol (Optiprep; Cat#1114542, Cosmo Bio) (40% (w/v) final concentration of iodixanol). A discontinuous iodixanol gradient was prepared as following: sample/iodixanol was applied to the bottom of the tube, followed by 35%, 30%, 25%, and 20% (w/v) iodixanol/PBS, covered by 15% (w/v) iodixanol/PBS. The density gradient centrifugation was performed at 141,000 g and 4 °C for 20 h in a SW28 swing rotor (Beckman Coulter). A total of 18 fractions were collected starting from the top.

### Screenings

HEK293T cells were transfected with each plasmid expressing an EBV gene using Lipofectamine 2000 (Thermo Fisher Scientific). On the following day, media were changed to DMEM supplemented with 5% exosome-depleted FBS (System Biosciences, Palo Alto, CA, USA). After 48 h, exosomes were purified from the conditioned media. To identify exosomal proteins, we analyzed purified exosomes by immunoblot. For analyzing modulators of infectivity, EBV-negative Akata(-) cells were treated with purified exosomes in RPMI1640 supplemented with 5% exosome-depleted FBS (System Biosciences, Palo Alto, CA, USA). After 6 h, cells were infected with EGFP-EBV. At 48 h after infection, cells were fixed with 4% paraformaldehyde. The infectivity was determined using a flow cytometer (FACS Canto, BD Biosciences, San Jose, CA, USA).

### Luciferase reporter assay

HEK293T cells were transfected with AP-1 reporter plasmid (pAP1-Luc) and promoterless-RLuc as an internal control using Lipofectamine 2000 (Thermo Fisher Scientific) and incubated in DMEM supplemented with 5% exosome-depleted FBS. After 48 h, cells were treated with purified exosome. Cells were lysed 24 h later and subjected to luciferase assays using the Dual-Luciferase Reporter Assay System (Promega).

### RNA-seq

Akata(-) cells were pre-treated with BGLF2-containing or control exosome for 16 h, and then infected with BGLF2-KO EBV at room temperature for 2 h with agitation. After infection, cells were resuspended in medium containing exosomes. The top 30,000 EBV-infected cells expressing EGFP were sorted by a FACS Aria II Cell Sorter (BD Biosciences) at 24 hpi.

Total RNA was extracted using the RNeasy Mini kit (Qiagen). Evaluation of RNA, RNA-seq library preparation, Illumina sequencing, and data preprocessing were performed as described previously (Okuno et al., 2019).

### qPCR

Total RNA was reverse-transcribed to cDNA using the PrimeScript II Reverse transcriptase kit (Takara Bio) or the *SuperPrep* II Cell Lysis & RT kit for qPCR (TOYOBO, Osaka, Japan). Viral DNA and mRNA levels were analyzed by qPCR using the 7500 Fast DX Real-Time PCR system (Applied Biosystems, Foster City, CA, USA) as previously described (Sato et al., 2017). The primer sequences used in this study are listed in Table 1.

**Table 1:**
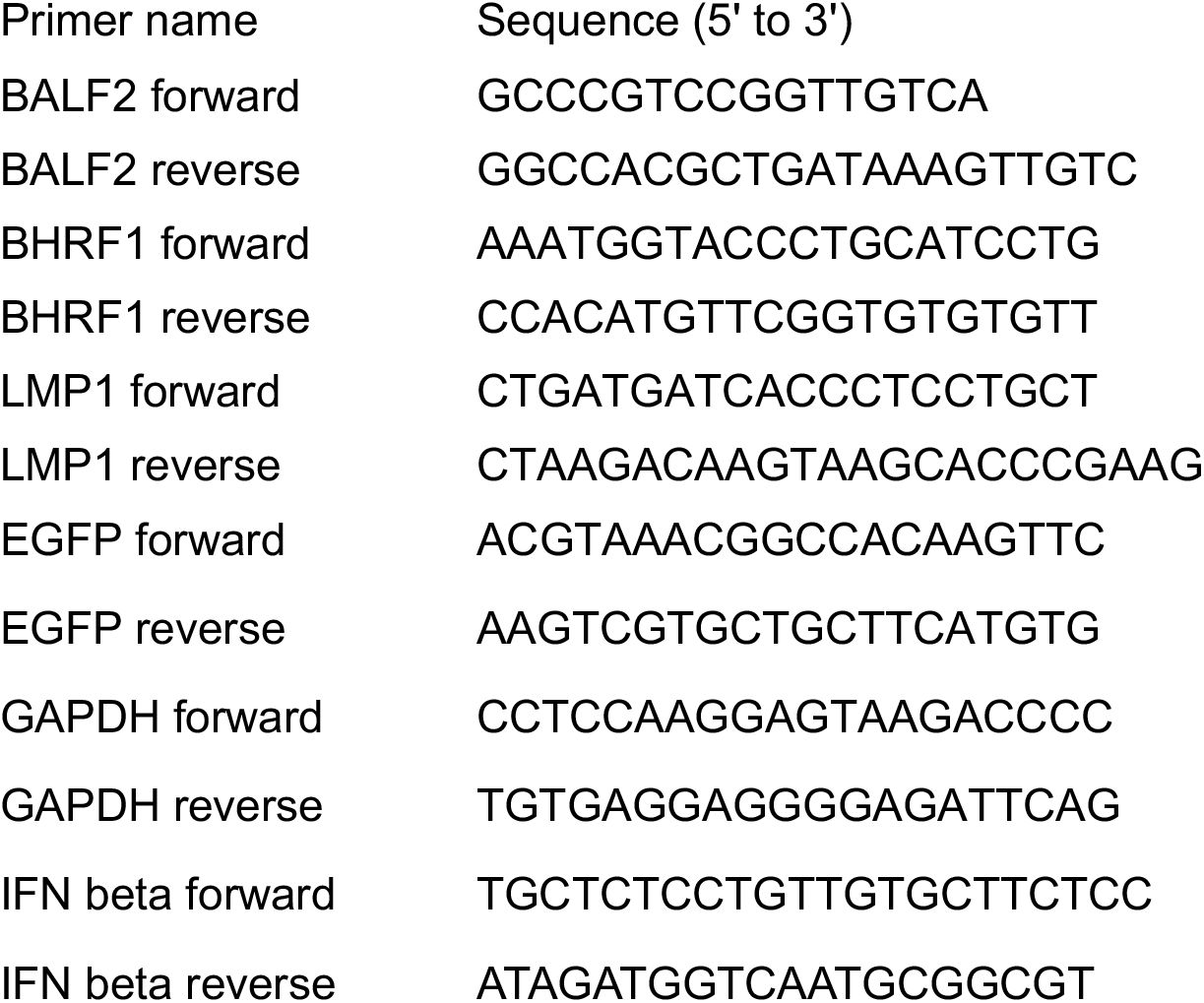
Oligonucleotide primers used for Qpcr.

### Statistical analysis

Results are shown as the means ± standard error (SE) of at least three independent experiments. Statistical analyses were performed using Microsoft Excel. Welch’s t-test was used to determine significance, and *P* < 0.05 was considered to be statistically significant.

## Data availability

All RNA-seq datasets have been deposited to the DNA Data Bank of Japan (DDBJ; https://www.ddbj.nig.ac.jp/index-e.html) under the accession number: DRA010388.

## Acknowledgements

The authors thank Hironori Yoshiyama for providing invaluable materials; Yohei Yamauchi for comments on the manuscript; Daisuke Yuasa, Shingo Ochiai, Shuko Kumagai, and Tomoko Kunogi for technical supports; Minoru Tanaka and the Division for Medical Research Engineering, Nagoya University Graduate School of Medicine for FACS analysis.

This work was supported in part by grants from the Japan Society for the Promotion of Science (JSPS) KAKENHI (https://www.jsps.go.jp) (Grant Numbers JP19H04829, JP21K15448 to Y.S., and JP20H03493 to H.K.); the JST (https://www.jst.go.jp) PRESTO (Grant Number JPMJPR19H5) to Y.S.; the Japan Agency for Medical Research and Development (AMED, https://www.amed.go.jp) (JP19fm0208016 and JP20wm0325012 to T.M., JP19ck0106517 to Y.O., and JP19jk0210023 to Y.S.); the Takeda Science Foundation (https://www.takeda-sci.or.jp) to Y.S. and T.M.; the Hori Sciences and Arts Foundation (https://www.hori-foundation.or.jp) to Y.S. and H.K.; the MSD Life Science Foundation (https://www.msd-life-science-foundation.or.jp) to Y.S.; and the Uehara Memorial Foundation (https://www.ueharazaidan.or.jp/) to H.K.. H.I. is supported by the JSPS Research fellowship (19J23589). T.S. and T.I. are supported by the Takeda Science Foundation scholarship.

## Figure Legends

**Supplementary Figure S1:**
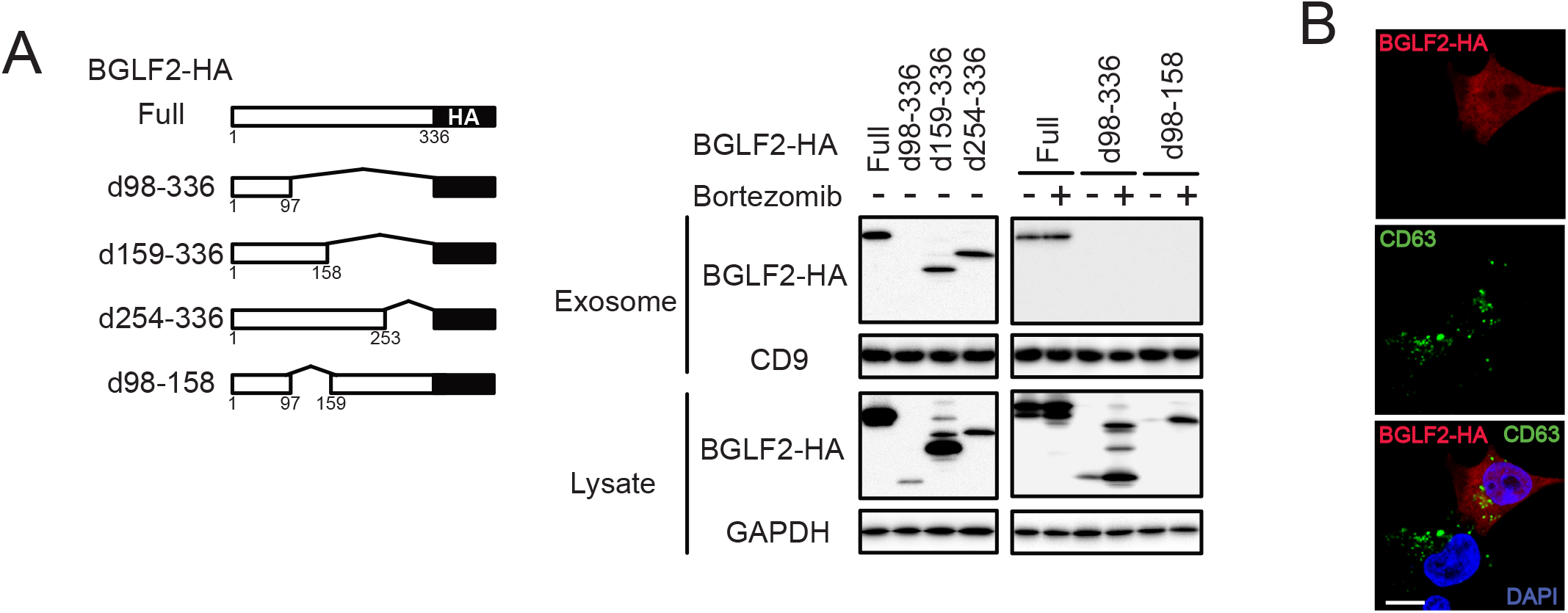
The internal region of BGLF2 is required for packaging into exosomes. (A) The C-terminal domain of BGLF2 was dispensable for incorporation into exosomes. HEK293 cells were transfected with a deletion mutant of BGLF2-expression plasmids and treated with or without 250 nM of Bortezomib for 24 h before harvest. Exosomes were isolated from the culture supernatant. (B) Cellular localization of BGLF2 and CD63 in HEK293 cells. HEK293 cells were transfected with HA-tagged BGLF2-expression plasmid. Cells were fixed at 2 days post-transfection and then stained with anti-HA and anti-CD63 antibodies.

**Supplementary Figure S2:**
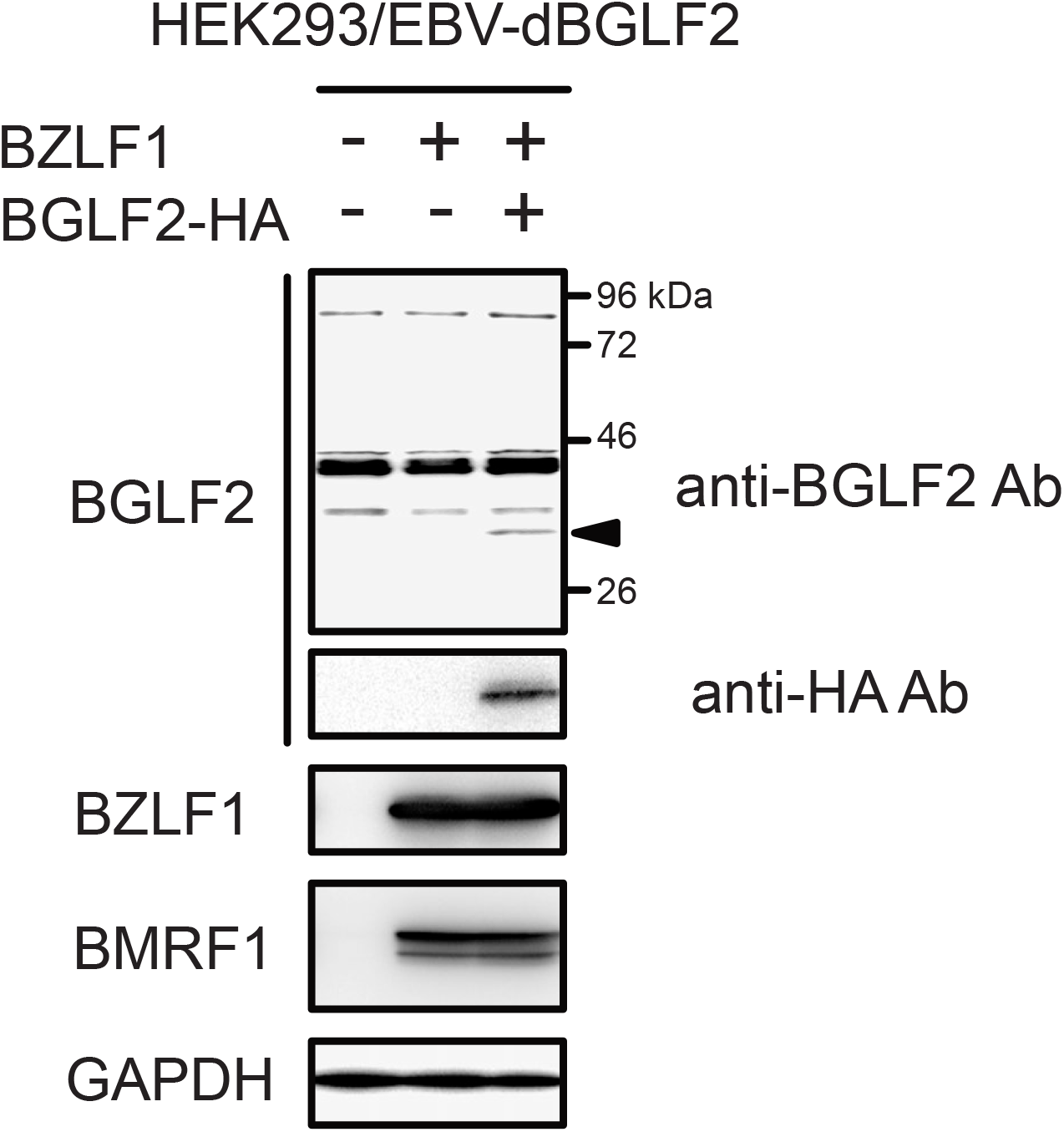
Detection of BGLF2 protein by anti BGLF2 antibody. HEK293/EBV-dBGLF2 cells were transfected with the BZLF1- and HA-tagged BGLF2- expression plasmids. Cells were harvested after 2 days and subjected to Western blotting with anti-BGLF2, anti-BZLF1, anti-BMRF1, and anti-GAPDH antibodies.

